# Partial reprogramming induces a steady decline in epigenetic age before loss of somatic identity

**DOI:** 10.1101/292680

**Authors:** Nelly Olova, Daniel J Simpson, Riccardo Marioni, Tamir Chandra

**Author notes:** to whom correspondence should be addressed: Dr Tamir Chandra, MRC Human Genetics Unit, Institute of Genetics & Molecular Medicine, University of Edinburgh EH4 2XU. These authors contributed equally to this work (Nelly Olova, Daniel J Simpson). Non-corresponding.

## Abstract

Induced pluripotent stem cells (IPSCs), with their unlimited regenerative capacity, carry the promise for tissue replacement to counter age-related decline. However, attempts to realise *in vivo* iPSC have invariably resulted in the formation of teratomas. Partial reprogramming in prematurely aged mice has shown promising results in alleviating age-related symptoms without teratoma formation. Does partial reprogramming lead to rejuvenation (i.e. “younger” cells), rather than dedifferentiation, which bears the risk of cancer? Here we analyse the dynamics of cellular age during human iPSC reprogramming and find that partial reprogramming leads to a reduction in the epigenetic age of cells. We also find that the loss of somatic gene expression and epigenetic age follow different kinetics, suggesting that they can be uncoupled and there could be a safe window where rejuvenation can be achieved with a minimised risk of cancer.

## Introduction, Results, Discussion

The human ageing process is accompanied by multiple degenerative diseases. Our understanding of such ageing related disorders is, nevertheless, fragmented, and the existence and nature of a general underlying cause are still much debated (Faragher 2015; Gladyshev & Gladyshev 2016). The generation of induced pluripotent stem cells (iPSCs) allows the reprogramming of somatic cells back to an embryonic stem cell (ESC) like state with an unlimited regenerative capacity. This has led to multiple strategies for tissue replacement in degenerative diseases (Takahashi et al. 2007). Clinical application of iPSCs however, is at its infancy (V. K. Singh et al., 2015; Soria-Valles et al., 2015; Takahashi & Yamanaka, 2016), and the potency of iPSCs bears risks, not least cancer induction. For example, *in vivo* experiments with iPSCs have shown that continuous expression of Yamanaka factors (Oct4, Sox2, Klf4 and c-Myc, thus OSKM) in adult mice invariably leads to cancer (Abad et al. 2013; Ohnishi et al. 2014).

To avoid this risk, a parallel concept of epigenetic rejuvenation has been proposed: the ageing process in cells can be reversed whilst avoiding dedifferentiation (Singh & Zacouto 2010; Manukyan & Singh 2012). In other words, an old dysfunctional heart cell could be rejuvenated without the need for it to be passed through an embryonic/iPSC state. The concept of epigenetic rejuvenation requires that rejuvenation and dedifferentiation each follow a distinct pathway. Nevertheless, it is not well understood whether rejuvenation and dedifferentiation are invariably intertwined, or instead whether it is possible to manipulate age without risking dedifferentiation.

The epigenetic rejuvenation potential of partial reprogramming with OSKM factors was previously shown by the forced expression of OSKM+*LIN28* in senescent human fibroblasts, which led to recovering the high mobility of histone protein 1β by day 9, a feature characteristic for young fibroblasts (Manukyan & Singh 2014). Ocampo et al. further demonstrated that partial reprogramming by transient cyclic induction of OSKM ameliorates signs of ageing and extends lifespan in progeroid mice, with no resulting teratoma formation (Ocampo et al. 2016). This established partial reprogramming as a promising candidate intervention for age-related disease. Estimating epigenetic age, which is currently the most promising proxy for biological age (Jylhävä et al. 2017; Wagner 2017), was, however, not possible to measure in mice at the time of the Ocampo study. This has left the nature (i.e. dedifferentiation/rejuvenation) of the described cellular changes unexplored:

1. Does the epigenetic remodelling seen truly reflect rejuvenation (i.e. a reduction in cellular/tissue age)? If so, can we observe a decrease in epigenetic age in partially reprogrammed human cells?
2. What is the extent of rejuvenation upon reaching a partially reprogrammed state (e.g. years of epigenetic age decrease)?
3. What are the dynamics of dedifferentiation in early reprogramming?

A major obstacle in understanding the relation between differentiation and ageing has been our inability to accurately measure cellular age with a high correlation to the chronological age of the organism. However, over the last five years a number of age predictors have been developed, the most accurate of which utilise DNA methylation (known as epigenetic clocks) (Horvath 2013; Hannum et al. 2013; Weidner et al. 2014; Levine et al. 2018; Horvath et al. 2018), with the first Horvath multi-tissue age-predictor being the most widely applicable and used (r=0.96). This “Horvath clock” shows the highest correlation to chronological age, predicting the age (or epigenetic age, eAge) of multiple tissues with a median error of 3.6 years (Horvath 2013). eAge is distinct from and poorly correlated with other age-related biomarkers, such as senescence and telomere length, which have been shown to correlate independently with the process of ageing (Lowe et al. 2016; Marioni et al. 2016). Moreover, an acceleration of epigenetic age as measured by the “Horvath clock” is associated with a higher risk of all-cause mortality (Marioni et al. 2015; Christiansen et al. 2016; Perna et al. 2016), premature ageing syndromes (Down and Werner) (Maierhofer et al. 2017; Horvath et al. 2015), frailty and menopause (Breitling et al. 2016; Levine et al. 2016). All of these studies suggest that eAge may capture a degree of biological ageing.

To understand the dynamics of eAge during reprogramming, we applied Horvath’s multi-tissue age predictor over a previously published reprogramming time-course on human dermal fibroblasts (HDFs) (Ohnuki et al. 2014; Horvath 2013). After OSKM transfection, successfully transformed subpopulations were isolated and analysed at regular time points during 49-days for gene expression and DNA methylation (detailed schematic shown in Supplementary Figure 1). Epigenetic rejuvenation, i.e. decrease of eAge, commenced between days 3 and 7 after OSKM transduction in the partially reprogrammed TRA-1-60 (+) cells (characterised in Tanabe et al. 2013) and continued steadily until day 20, when eAge was stably reset to zero (Fig. 1a). A broken stick model (comprising two linear regressions joined at a break-point), showed a good fit to the observed data starting from day 3, and measured a steady decrease with 3.8 years per day until day 20 (SE 0.27, P = 3.8×10^-7^) (Fig. 1a). The TRA-1-60 (+) cell populations at days 7 and 11 have been previously characterised as ‘partially reprogrammed’ for their high expression of pluripotency markers but also high reversion rates towards somatic state (Tanabe et al. 2013). Therefore, the observed eAge decline at days 7 and 11 suggests that partial reprogramming can indeed be considered a rejuvenation mechanism in human cells.

**Figure 1.**
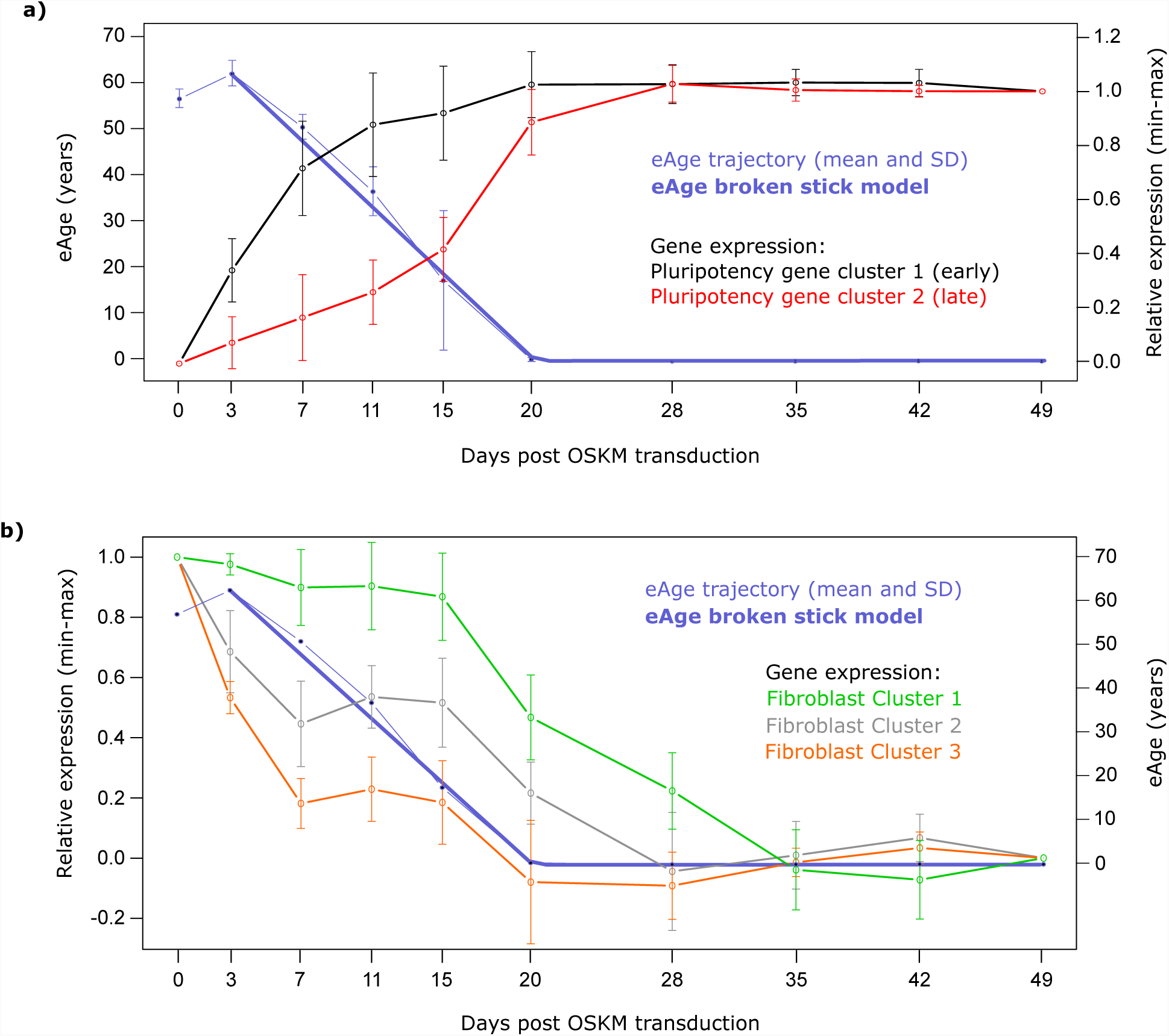
Dynamics of eAge and gene expression in a 49-day HDF reprogramming time-course. **(a) Left Y axis**: eAge trajectory of Horvath’s multi-tissue age predictor calculated from DNA methylation arrays from the following cell populations: day 0 (HDFs), day 3 (OSKM-expressing EGFP (+) HDFs), day 7, 11, 15, 20 and 28 (human pluripotency marker TRA-1-60 (+) cells at intermediate stages of reprogramming), and fully reprogrammed iPSCs from days 35, 42 and 49. Data was fit with a broken stick model composed of two linear sections. Error bars represent SD. Measured rate (years per day) of eAge decrease [day 3 - day 20] = −3.8, SE 0.27, P = 3.8 x 10^-7^. **Right Y axis**: Composite gene expression trajectories of key pluripotency markers, statistically clustered as per Genolini et al. 2016. Microarray expression data was obtained for the same time points and cell populations as for eAge. Relative expression values were LOG2 transformed and presented as arbitrary units starting from ‘0’ for ‘day 0’ to ‘1’ for ‘day 49’. Error bars represent SD. **(b) Left Y axis**: Composite gene expression trajectories of key fibroblast markers statistically clustered as described for the pluripotency markers in (a). Relative expression values are presented as arbitrary units starting from ‘1’ for ‘day 0’ to ‘0’ for ‘day 49’. **Right Y axis:** eAge as in (a) left Y axis, without SD.

Horvath’s multi-tissue age predictor is the most accurate and widely used for various cell types and tissues (Wagner 2017). Nevertheless, we calculated eAge from alternative DNA methylation-based age predictors: four tissue-specific clocks (Hannum et al. 2013; Weidner et al. 2014; Horvath et al. 2018), one that incorporates clinical measures, called PhenoAge (Levine et al. 2018), and individual CpGs previously correlated with age (Garagnani et al. 2012). All clocks consistently reached the point of reset to their iPSC eAge at day 20, despite the cells not being fully reprogrammed before day 28 (Ohnuki et al. 2014) (Supplementary Figure 2). Again, eAge showed a steady decline from day 3 to day 20 in the skin & blood and Weidner 99 CpG clocks, PhenoAge declined from day 7 to day 20, while the Hannum and Weidner 3 CpG clocks did not produce informative trajectories. Overall, eAge values and ‘years’ of decrease varied between the clocks (actual chronological age of HDF donors is not available for reference) (Supplementary Figure 2). The highest age associated individual CpG (*ELOVL2*’s cg16867657) showed a similar trajectory to the Horvath eAge decline, however, the remaining CpGs produced inconsistent trajectories (Supplementary Figure 2). The observed differences are not surprising, given the alternative clocks were validated for blood (Hannum et al. 2013; Weidner et al. 2014), forensic applications (Horvath et al. 2018), whole organisms (Levine et al. 2018) or various tissues as for the individual CpGs (Garagnani et al. 2012).

In Ocampo et al. partial reprogramming was achieved after just two days of OKSM induction in mice carrying an inducible OSKM transgene (Ocampo et al. 2016). However, such ‘secondary’ systems for direct reprogramming are known to have up to 50-fold higher efficiency and accelerated kinetics in comparison to virally transduced *in vitro* systems (Wernig et al. 2008). To facilitate comparison to other systems and associate eAge with intermediate states in the reprogramming trajectory we compared it to gene expression measured in the same samples. We analysed corresponding microarray expression data for 19 well-established pluripotency marker genes (Table 1 and Supplementary fig.3) as a proxy for reaching a mature pluripotent state (Ginis et al. 2004; Cai et al. 2006; Mallon et al. 2013; Galan et al. 2013; Boyer et al. 2005). We statistically clustered the expression patterns of those genes (Genolini et al. 2015), which resulted in two composite trajectories. These followed previously described expression dynamics of early (cluster 1) and late (cluster 2) activated pluripotency genes (Fig. 1a) (Tanabe et al. 2013; Chung et al. 2014; Buganim et al. 2012; Takahashi & Yamanaka 2016). Pluripotency gene cluster 1 included *NANOG, SALL4, ZFP42, TRA-1-60, UTF1, DPPA4* and *LEFTY2*, and their expression increased dramatically within the first 10 days and then established stable pluripotency expression levels by day 20. In contrast, pluripotency gene cluster 2 (containing late expressing genes such as *LIN28, ZIC3* and *DNMT3B*) elevated expression more slowly and reached stable pluripotency levels by day 28 (Tanabe et al. 2013; Chung et al. 2014). Interestingly, eAge reset to zero at the same time that the genes in cluster 1 reached their pluripotent state levels, which temporally precedes full pluripotency. This also coincided with a peak in expression of a number of embryonic developmental genes between days 15 and 20, and might suggest that the reset marks a point where the cells reach an embryonic-like state but are not yet fully pluripotent (Table 1 and Supplementary Figure 4). In summary, eAge decline is observed well within the first wave of pluripotency gene expression.

**Table 1.** List of pluripotency and fibroblast marker genes used in gene expression clusters. Key pluripotent marker genes were selected from Ginis et al. 2004; Cai et al. 2006; Mallon et al. 2013; Galan et al. 2013; Boyer et al. 2005. Fibroblast marker genes were selected from Kalluri & Zeisberg 2006; Zhou et al. 2016; Janmaat et al. 2015; Pilling et al. 2009; Chang et al. 2014; Goodpaster et al. 2008; MacFadyen et al. 2005.

| Marker | Gene | Protein name | Accession | Cluster |
| --- | --- | --- | --- | --- |
| Pluripotency | *NANOG* | Nanog homeobox | A_23_P204640 | 1 (early) |
| Pluripotency | *REX1 (ZFP42)* | Zinc Finger Protein 42 | A_23_P395582 | 1 (early) |
| Pluripotency | *TRA-1-60/81 (PODXL)* | Podocalyxin | A_23_P215060 | 1 (early) |
| Pluripotency | *UTF1* | Undifferentiated embryonic cell transcription factor 1 | A_33_P3294217 | 1 (early) |
| Pluripotency | *DPPA4* | Developmental pluripotency associated 4 | A_23_P380526 | 1 (early) |
| Pluripotency | *TDGF1 (CRIPTO)* | Teratocarcinoma-derived growth factor 1 | A_23_P366376 | 1 (early) |
| Pluripotency | *SALL4* | Spalt like transcription factor 4 | A_23_P109072 | 1 (early) |
| Pluripotency | *LEFTY1* | Left-right determination factor 1 | A_23_P160336 | 1 (early) |
| Pluripotency | *LEFTY2* | Left-right determination factor 2 | A_23_P137573 | 1 (early) |
| Pluripotency | *DNMT3A* | DNA methyl-transferase 3A | A_23_P154500 | 1 (early) |
| Pluripotency | *TFCP2L1* | Transcription factor CP2 like 1 | A_23_P5301 | 1 (early) |
| Pluripotency | *TERF1* | Telomeric repeat binding factor (NIMA-interacting) 1 | A_23_P216149 | 2 (late) |
| Pluripotency | *DPPA5* | Developmental pluripotency associated 5 | A_32_P233950 | 2 (late) |
| Pluripotency | *TERT* | Telomerase reverse transcriptase | A_23_P110851 | 2 (late) |
| Pluripotency | *ZIC3* | Zic family member 3 | A_23_P327910 | 2 (late) |
| Pluripotency | *LIN28a* | LIN28 homolog A | A_23_P74895 | 2 (late) |
| Pluripotency | *LIN28b* | LIN28 homolog B | A_33_P3220615 | 2 (late) |
| Pluripotency | *LECT1* | Leukocyte cell derived chemotaxin 1 | A_23_P25587 | 2 (late) |
| Pluripotency | *DNMT3B* | DNA methyl-transferase 3B | A_23_P28953 | 2 (late) |
| Fibroblast | *COL3A1* | Pro-collagen α2(III) | A_24_P935491 | 1 |
| Fibroblast | *FSP-1* | Fibroblast surface protein | A_23_P94800 | 1 |
| Fibroblast | *TGFB3* | Transforming growth factor beta 3 | A_23_P88404 | 1 |
| Fibroblast | *TGFB2* | Transforming growth factor beta 2 | A_24_P402438 | 1 |
| Fibroblast | *COL1A2* | Pro-collagen α2(I) | A_24_P277934 | 2 |
| Fibroblast | *ITGA1* | Integrin α1b1 (VLA-1) | A_33_P3353791 | 2 |
| Fibroblast | *DDR2* | Discoidin-domain-receptor-2 | A_23_P452 | 2 |
| Fibroblast | *P4HA3* | Prolyl 4-hydroxylase | A_24_P290286 | 2 |
| Fibroblast | *THY1* | Thy-1 cell surface antigen; CD90 | A_33_P3280845 | 2 |
| Fibroblast | *FAP* | Fibroblast activation protein | A_23_P56746 | 2 |
| Fibroblast | *CD248* | Endosialin, TEM1 | A_33_P3337485 | 2 |
| Fibroblast | *VIM* | Vimentin | A_23_P161190 | 2 |
| Fibroblast | *COL1A1* | Pro-collagen α1(I) | A_33_P3304668 | 3 |
| Fibroblast | *ITGA5* | Integrin α5b1 | A_23_P36562 | 3 |
| Fibroblast | *P4HA1* | Prolyl 4-hydroxylase | A_33_P3214481 | 3 |
| Fibroblast | *P4HA2* | Prolyl 4-hydroxylase | A_33_P3394933 | 3 |
| Fibroblast | *TGFB1* | Transforming growth factor beta 1 | A_24_P79054 | 3 |
| Fibroblast | *HSP47* | Serpin family H member 1, SERPINH1 | A_33_P3269203 | - |
| Fibroblast | *CD34* | Hematopoietic progenitor cell antigen | A_23_P23829 | - |

Therapeutic partial reprogramming will depend on rejuvenation with minimal dedifferentiation, which carries the risk of malignancies. We studied the dynamics of fibroblast gene down-regulation as a proxy for the loss of somatic cell identity. The individual trajectories of 19 commonly used fibroblast marker genes (Kalluri & Zeisberg 2006; Zhou et al. 2016; Janmaat et al. 2015; Pilling et al. 2009; Chang et al. 2014; Goodpaster et al. 2008; MacFadyen et al. 2005) (Table 1 and Supplementary Fig. 5) clustered into three composite expression patterns, two of which (clusters 2 and 3) went into an immediate decline after OSKM induction (Fig. 1b). However, one fibroblast-specific cluster (cluster 1) remained stable in its expression for the first 15 days. Interestingly, after day 7, fibroblast-specific gene expression in clusters 2 and 3 stopped declining and plateaued until day 15, coinciding with a peak in expression of senescence markers between days 11 and 15 (Supplementary Figure 6). Vimentin (*VIM*), for example, remained at 60% of maximal expression until day 15 of reprogramming, similarly to *FAP, CD248* and *COL1A2* in cluster 2 (Supplementary fig. 5). After day 15, fibroblast gene expression declined rapidly in all three clusters, and only by day 35 had all reached ESC expression levels, marking a complete loss of somatic identity (Fig. 1b). Cluster 1, which contains the well described indicators of fibroblast identity *FSP1, COL3A1* and *TGFB2/3* (Kalluri & Zeisberg 2006), showed the slowest decline, and was also the last to reach ESC expression levels. In summary, we found that a number of fibroblast specific genes maintained high expression levels until day 15, by which time a substantial drop in eAge has been observed.

Epigenetic rejuvenation or the reversal of cellular age, is a promising concept as it could avoid the oncogenic risks associated with dedifferentiation. Here, we analysed a reprogramming time-course on HDFs and show that eAge declines in partially reprogrammed cells before their somatic identity is entirely lost.

It is well established that partial reprogramming happens within an early, reversible phase during the iPSC reprogramming time-course, which involves the stochastic activation of pluripotency genes. It is followed by a more deterministic maturation phase with predictable order of gene expression changes, where cell fate is firmly bound towards pluripotency (Takahashi & Yamanaka 2016; Smith et al. 2016). Indeed, it has been shown that mouse fibroblasts fail to become iPSC and revert to their original somatic state if OSKM expression is discontinued during the initial stochastic phase (Brambrink et al. 2008; Stadtfeld et al. 2008). Previously, Tanabe et al. showed that TRA-1-60 (+) cells at reprogramming days 7 and 11 have not yet reached maturation and are partially reprogrammed (Tanabe et al. 2013) but our analysis already shows a decrease in their eAge according to multiple age predictors (Fig. 1a and Supplementary Figure 2). We have also shown that a large proportion of fibroblast marker genes maintain relatively high levels of expression until day 15 (Fig. 1b and Supplementary Figure 5). Nearly unchanged levels of expression on day 15 were previously also shown for a large proportion of somatic genes (Tanabe et al. 2013). Together with increased senescence gene expression between days 11 and 15 (Supplementary Figure 6), this likely contributes to the high propensity of partially reprogrammed TRA-1-60 (+) cells to revert back to somatic phenotype before day 15 in the time-course (Tanabe et al. 2013). Interestingly, the step-wise decline of fibroblast gene expression coinciding with a peak in expression of senescence genes seems to delay the loss of somatic identity but not the expression of pluripotency genes. Taken together, the different dynamics between the step-wise fibroblast expression and the linear decline in eAge further indicate that dedifferentiation and epigenetic rejuvenation can be uncoupled.

Our data suggest a window of opportunity within the uncommitted reprogramming phase, where a decline of eAge happens alongside partial maintenance of fibroblast gene expression. A deeper understanding of the kinetics of rejuvenation will be required to master therapeutic partial reprogramming, since any progress of dedifferentiation, even in a small subpopulation, carries the risk of malignancies. Our bulk expression analysis does not allow for a precise definition of the safe rejuvenation boundaries, and further experiments on a single cell level and in *in vivo* conditions are needed to determine a safe epigenetic rejuvenation window in different reprogramming systems. Upon defining safe boundaries, consideration should also be given to the steep decline of eAge, which resets to zero well ahead of the establishment of a pluripotent state, according to a number of age predictors (Supplementary Figure 2). Most likely this marks the point of reaching prenatal or embryonic stage, as suggested by the peak in expression of key developmental genes (Supplementary Figure 4).

The extent of epigenetic rejuvenation in years (human) or months (mouse), which can be achieved through partial reprogramming, also needs further attention and will most likely differ with the different reprogramming systems. The ‘Horvath clock’ shows up to 10 years of rejuvenation in Ohnuki et al.’s system by day 7 and another 10+ years by day 11. However, the intrinsic median estimation error of 3.6 years in this age predictor, the varying eAge rejuvenation values between the different age predictors, and the intra-replicate biological variation seen from the large error bars, highlight the need for more experiments and repetitions before this is established with a higher certainty.

Despite the obvious differences in reprogramming kinetics, our results also suggest that the improvements observed by Ocampo et al. in their OSKM-inducible secondary reprogramming system, might be due to epigenetic rejuvenation. It remains to be shown how stable in time the rejuvenated phenotype is in either of the systems. Further analysis is also needed regarding the effect of partial reprogramming on adult stem cells or premalignant cells, which have already shown a higher propensity of transforming to malignancy (Abad et al. 2013; Ohnishi et al. 2014). It is possible that a premalignant phenotype could be attenuated or amplified by partial reprogramming. In summary, our findings reveal exciting possibilities but also open a number of questions and highlight areas that need further attention.

## Acknowledgements

We thank Chris Ponting, Steve Horvath and Keisuke Kaji for their helpful advice and comments on the manuscript.

## Conflict of interest

The authors of this paper have no conflicts of interest to declare.

## Supplemental Experimental procedures

### Overview of the Ohnuki et al. experimental setup and datasets

450K DNA methylation array and gene expression microarray data of full HDF reprogramming time-course was obtained from GSE54848. A schematic of experimental setup and time points is provided in Supplementary Figure 1. Briefly, HDF cells were transfected with EGFP-labelled OSKM on day 0 and cultured in virus-containing medium for 24 hours, then replaced by 10% FBS-containing medium for 8 days before replacing with human ESC medium. EGFP (+) cells, representing the population of successfully transfected cells, which permanently express the OSKM factors, were sorted by flow cytometry on day 3. Intermediate reprogrammed cells positive for the pluripotency marker TRA-1-60 were sorted by magnetic activated cell sorting on days 7, 11, 15, 20 and 28 post-transfection. Day 28-sorted TRA-1-60 (+) cells were further expanded and samples collected three more times on each seventh day, i.e. on days 35, 42 and 49. Thus, sorted and collected cells at each time point were subjected to both gene expression and DNA methylation array analysis. Microarray gene expression (data available as LOG2 transformed) was performed for three to four replicates per data point, whilst DNA methylation data was performed for two to three replicates per time point.

### Predicting eAge

The pre-processed 450K DNA methylation array matrix of average methylation per CpG site of the full HDF reprogramming time-course was obtained from GSE54848 (downloaded using getGEO function from GEOquery package) and uploaded to the online DNA methylation age calculator to assess eAge: https://labs.genetics.ucla.edu/horvath/dnamage/ (Horvath 2013). Data processing including Horvath’s normalisation was performed according to tutorial guidelines. Missing CpG values were imputed by Horvath’s online DNAm age calculator. During QC, around 1600 CpGs were lost, therefore methylation data for each time point contained 26,987 CpG sites out of the suggested 28,587 CpGs, a fact unlikely to have any significant impact on the normalisation or age prediction. PhenoAge, skin & blood, Hannum, Weidner 99 and 3 CpG age predictors were applied to average methylation values. Missing CpG values were imputed as zero before applying these age predictors.

All ages presented in the manuscript are calculated eAges, no actual ages of HDF donors were available.

### Methylation Age Trajectories

For the Horvath multi-tissue age predictor, a ‘broken stick’ model with two linear sections was constructed to chart overall change in DNA methylation age over time between the three HDF cell lines. A linear mixed model was then specified with a random intercept term for each replicate. A variable break point was set between the minimum and maximum day, plus and minus a small constant (3 days), respectively. The predicted values from the regression models were plotted against the measurement day. For the all other age predictor plots (Supplementary Figure 2), mean eAge was calculated for all samples at each time point (2-3 samples depending on the time point) and plotted against time during the time-course. Standard deviation for eAge was also calculated and plotted as error bars at each time point.

### Gene clusters and trajectories

For each gene in a category (e.g. pluripotent gene list), a loess curve with a span of 0.5 was fitted with the predicted values extracted at each time point. The predicted values were then normalised within each gene to a value of 1 at the first time point and a value of 0 and the last time point (and vice versa for the pluripotent genes). K-means clustering for longitudinal data was applied to determine the optimal number of trajectories within each gene category.

All analyses were performed in R, using the kml (Genolini et al. 2015), lme4 (Bates et al. 2014), and lmerTest (Kuznetsova et al. 2016) packages.

